# Hybridization drives genetic erosion in sympatric desert fishes of western North America

**DOI:** 10.1101/644641

**Authors:** Tyler K. Chafin, Marlis R. Douglas, Bradley T. Martin, Michael E. Douglas

**Affiliations:** Department of Biological Sciences, University of Arkansas, Fayetteville, Arkansas 72701, USA

**Keywords:** climate change, ddRAD, hybridization, southwestern North America, species concepts

## Abstract

Many species have evolved or currently coexist in sympatry due to differential adaptation in a heterogeneous environment. However, anthropogenic habitat modifications can either disrupt reproductive barriers or obscure environmental conditions which underlie fitness gradients. In this study, we evaluated the potential for an anthropogenically-mediated shift in reproductive boundaries that separate two historically sympatric fish species (*Gila cypha* and *G. robusta*) endemic to the Colorado River Basin using ddRAD sequencing of 368 individuals. We first examined the integrity of reproductive isolation while in sympatry and allopatry, then characterized hybrid ancestries using genealogical assignment tests. We tested for localized erosion of reproductive isolation by comparing site-wise genomic clines against global patterns and identified a breakdown in the drainage-wide pattern of selection against interspecific heterozygotes. This, in turn, allowed for the formation of a hybrid swarm in one tributary, and asymmetric introgression where species co-occur. We also detected a weak but significant relationship between genetic purity and degree of consumptive water removal, suggesting a role for anthropogenic habitat modifications in undermining species boundaries. In addition, results from basin-wide genomic clines suggested that hybrids and parental forms are adaptively non-equivalent. If so, then a failure to manage for hybridization will exacerbate the long-term extinction risk in parental populations. These results reinforce the role of anthropogenic habitat modification in promoting interspecific introgression in sympatric species by relaxing divergent selection. This, in turn, underscores a broader role for hybridization in decreasing global biodiversity within rapidly deteriorating environments.

## Introduction

Many natural populations respond to anthropogenic change by either shifting geographic distributions or adjusting life histories so as to ‘track’ optimal conditions (Hoffmann and Sgrò 2011; Pecl *et al*. 2017). However, the ability of organisms to track changing environments is conditioned upon the rate of environmental change (Lindsey *et al*. 2013) and the rate at which adaptive machinery can act (Orr and Unckless, 2014). This evolutionary caveat creates an incentive for hybridization, in that recombinant genotypes might more rapidly establish in a dynamic adaptive landscape (Klonner *et al*. 2017). Widespread hybridization thus may provide an effective mechanism of population persistence in changing or novel conditions (Pease *et al*. 2016; Meier *et al*. 2017). Introgressed alleles which are beneficial under novel conditions can then be driven to fixation by the combined action of recombination and selection (Arnold and Martin 2010).

However, the relationship between hybridization and extinction is not well established under contemporary timescales. On one hand, hybrid lineages might facilitate adaptation by providing access to a greater pool of genetic variation (Dittrich-Reed and Fitzpatrick 2013; Schumer *et al*. 2018), whereas on the other, diversity might diminish as species boundaries dissolve (Buerkle *et al*. 2003; Kearns *et al*. 2018). Often, results are a combination of the above. Introgressed genotypes may initially compensate for erratic conditions and facilitate population persistence in the near term, but with lineages eventually merging if environmental change is prolonged (Seehausen *et al*. 2008). This presents an obvious paradox for the conservation biologist, in that the permeability of species-boundaries may be seen as promoting both persistence and extinction.

Hybridization also represents a legacy issue for conservation policy (Allendorf *et al*. 2001), due primarily to its conflict with a species-centric management paradigm (Fitzpatrick *et al*. 2015; Hamilton and Miller 2016). Although the reticulate nature of speciation has become a contemporary research focus (e.g. Mallet *et al*. 2016), it has yet to gain consensus among managers (vonHoldt *et al*. 2018). This unanimity is required to understand the manner by which anthropogenic modifications disrupt species boundaries (Grabenstein and Taylor 2018; Ryan *et al*. 2018). However, predicting the outcome of hybridization in a changing environment requires an understanding of both the temporal and spatial stability of the mechanisms (e.g. intrinsic and extrinsic) that are responsible for maintaining species boundaries. In this sense, consistent patterns can often be obscured by local context (e.g. individual behaviors, population demographics; Klein *et al*. 2017). Hence, there remains a need to quantify the manner by which species boundaries in diverse taxa respond to rapid environmental change. We applied these perspectives to endemic, large-bodied and long-lived minnows that exist within the Colorado River, one of the most impacted riverine ecosystems of the Anthropocene (Hughes *et al*. 2007). Because of the pervasive human impacts therein, the Colorado River provides a natural laboratory within which to examine the stability of species undergoing rapid, anthropogenically-induced environmental change.

### Hybridization in Gila

Hybridization has long been recognized as an evolutionary process in fishes (Hubbs, 1955), and as such, has been hypothesized as a mechanism for native fish diversification in western North America (e.g. DeMarais *et al*. 1992). External fertilization and limited predisposition for pre-mating isolation also likely promote hybridization as a common mechanism throughout fishes (e.g. Mayr 1963). An inseparable link also exists between fishes and their environment, such that opportunities for hybridization are controlled by characteristics of the riverscape (Hopken *et al*. 2013; Thomaz *et al*. 2016). The instability produced by modified flows may compromise boundaries between historically coexisting species, or provide ecological opportunities within which hybrid lineages might capitalize (Dowling and Secor 1997). The fact that habitats in western North America were historically subjected to tectonism and progressive aridity also provides an explanation for the high prevalence of hybridization relative to species richness in southwestern North America (e.g. Mandeville *et al*. 2017; Bangs *et al*. 2018). However, more contemporary anthropogenic modifications are also prominent and widespread, most apparent in the form of water acquisition and retention (Cayan *et al*. 2010). As a result, niche gradients that historically segregated species are now seriously perturbed. This, in turn, can promote hybridization by effectively removing selection against hybrid phenotypes, and by disrupting the phenology and reproductive cues that discourage heterospecific mating (Grabenstein and Taylor 2018).

We applied these perspectives to three species of conservation concern endemic to the Colorado River Basin: Humpback chub [*Gila cypha* (IUCN status=Endangered)], Roundtail chub [*G. robusta* (Near Threatened)], and Bonytail [*G. elegans* (Critically Endangered)]. All are hypothesized as exhibiting various levels of historic hybridization, with contemporary populations shaped by geologic processes and anthropogenic interventions. *Gila cypha* and *G. robusta*, display not only morphological intergradation (McElroy and Douglas 1995) but also taxonomic ambiguity (Douglas *et al*. 1989) and cannot be distinguished on the basis of mitochondrial (mt)DNA (Douglas and Douglas 2007; Dowling and DeMarais 1993), despite numerous lines of evidence supporting evolutionary independence [genetic structuring in nuclear markers (microsatellites: Douglas and Douglas 2007); discrete persistence in the fossil record (e.g. Uyeno and Miller 1963, 1965); pre-mating isolation in the form of exclusive reproductive ecology and phenology (Kaeding *et al*. 1990); and divergent phenotypic evolution (Smith *et al*. 1979; Valdez *et al*. 1990; Portz and Tyus 2004)]. Also, McElroy and Douglas (1995) and Douglas *et al*. (1998) found clear species-level differentiation in discriminant and geometric morphometric space, respectively, while the former also reported species-intermediacy at two sympatric localities (Desolation and Cataract canyons).

A parsimonious explanation for this mosaic pattern would invoke historic separation followed by contemporary hybridization. We tested this hypothesis herein and framed our results within the context of change both on geologic and contemporary timescales.

## Methods

### Sampling

Fin tissue was non-lethally sampled from 368 specimens across three native *Gila* of the Colorado River Basin (*G. cypha, G. elegans*, and *G. robusta*; Table 1), collected primarily by state/ federal agencies between 1997-2017 (see Acknowledgements). One location, at the San Rafael River, was sampled both in 2009 and 2017. Given the conservation status of these fishes, we minimized impacts on already-stressed populations by opportunistic sampling which took advantage of monitoring activities by agencies.

**Table 1:** Sampling locations for *Gila robusta, G. cypha* and *G. elegans*. Site=abbreviated locality identifier for each species, Major Drainage=River, Location=geographic site, County, State=per sampling site, and *N*=Number of samples excluding those that sequenced with sufficient coverage.

| Site | Major Drainage | Location | County, State | *N* |
| --- | --- | --- | --- | --- |
| *Gila robusta* |  |  |  |  |
| RBKR | Colorado | Black Rocks Canyon | Mesa, CO | 11 |
| RDES | Green | Desolation Canyon | Uintah, UT | 23 |
| RC15 | Colorado | 15-mile reach | Mesa, CO | 10 |
| RECC | Little Colorado | East Clear Creek | Coconino, AZ | 16 |
| RMCO | San Juan | Mancos River | Montezuma, CO | 10 |
| RMGR | Green | Middle Green R. tributaries | Sweetwater, WY | 24 |
| RNJA | San Juan | Navajo R. | Rio Arriba, NM | 10 |
| RLSR | Yampa | Little Snake R. tributaries | Carbon, WY | 31 |
| RLYC | Yampa | Little Yampa Canyon | Moffat, CO | 11 |
| RSRR | Green | San Rafael R. | Emery, UT | 23 |
| RUGR | Green | Upper Green R. tributaries | Sublette, WY | 16 |
| RWRW | White | White R. mainstem near Weaver Cn. | Uintah, UT | 15 |
| RWWC | Colorado | Colorado mainstem near Westwater Cn. | Grand, UT | 11 |
| RYAM | Yampa | Yampa R. mainstem | Moffat, CO | 15 |
| *Gila cypha* |  |  |  |  |
| HBKR | Colorado | Black Rocks Canyon | Mesa, CO | 18 |
| HDES | Green | Desolation Canyon | Uintah, UT | 24 |
| HGCN | Colorado | Grand Canyon | Coconino, AZ | 37 |
| HLCR | Little Colorado | Atomizer Falls and Colorado confluence | Coconino, AZ | 22 |
| HWWC | Colorado | Westwater Canyon | Grand, UT | 22 |
| HYAM | Yampa | Yampa Canyon | Moffat, CO | 8 |
| *Gila elegans* | Hatchery stock | USFWS Hatchery at Dexter, NM | Chaves, NM | 11 |

*Gila cypha* is constrained within five known aggregates associated with specific geomorphic features: Black Rocks, Cataract, Desolation, Grand, Westwater, and Yampa canyons (Fig. 1; USFWS, 2011), all of which were sampled save Cataract Canyon. Westwater and Black Rocks were treated separately, despite their potential for connectivity (Francis *et al*. 2016). Due to its range-wide extirpation, samples of *G. elegans* were obtained from the Southwestern Native Aquatic Resources and Recovery Center, Dexter, NM (formerly the Dexter National Fish Hatchery). Our sampling of *G. robusta* encompassed its entire range, to include pre-defined MUs (=Management Units; Douglas and Douglas 2007) and represented wild populations, with the exception the Mancos River, which was obtained from the Colorado Department of Wildlife Native Aquatic Species Restoration Facility. *Gila robusta* from the lower basin Bill Williams and Gila River drainages was not included, given its known polyphyly (Dowling and DeMarais 1993; Chafin *et al*. unpubl.).

**Figure 1:**
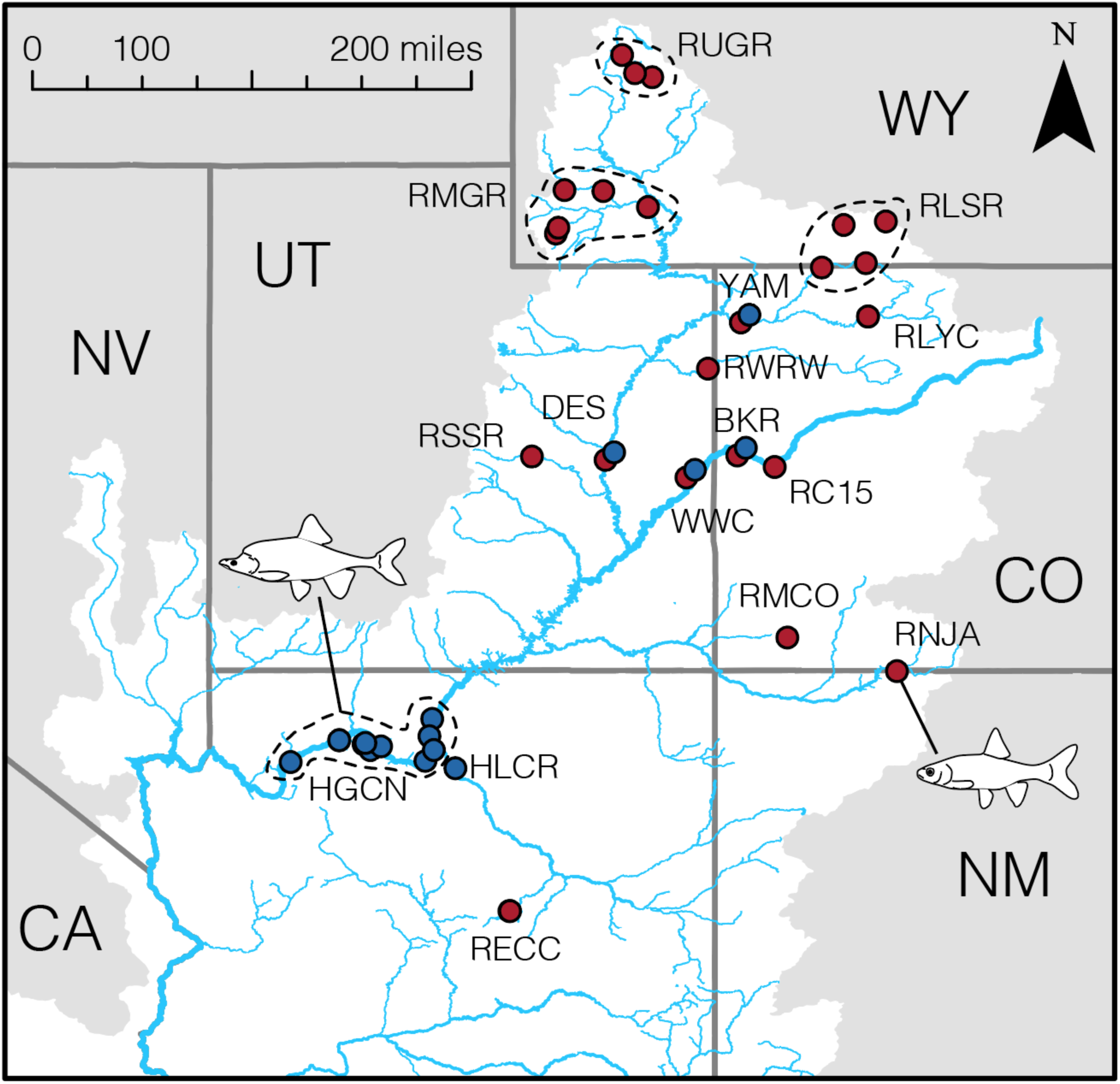
Sampling localities for *Gila cypha* (blue) and *G. robusta* (red) within the Colorado River Basin, western North America. Locality codes are defined in Table 2. Sympatric locations (BKR, DES, WWC, YAM) are slightly offset for visibility purposes. Inset cartoons the respective morphologies of each species.

### Data collection

Genomic DNA was extracted using either PureGene® or DNeasy® kits (Qiagen Inc.), with electrophoresis (2% agarose gel) confirming presence of sufficiently high molecular weight DNA. Our ddRAD library preparations were modified from previous protocols (Peterson *et al*. 2012). Restriction enzyme pairings and size-selection ranges were optimized using an *in silico* procedure (FRAGMATIC; Chafin *et al*. 2018). Samples were digested with *Msp*I (5’-CCGG-3’) and *Pst*I (5’-CTGCAG-3’) following manufacturer’s protocols (New England Biosciences). Fragments were then purified using Ampure XP beads (Beckman-Coulter Inc.) and concentrations standardized at 100ng per sample. Custom adapters containing in-line barcodes were ligated with T4 Ligase (New England Biosciences), pooled in sets of 48, and size-selected with the Pippin Prep (Sage Sciences) at 250-350bp prior to adjusting for adaptor length. We then utilized a 12-cycle PCR to extend adapters with indexed Tru-Seq primers and Phusion high-fidelity DNA polymerase (manufacturer protocols; New England Biosciences). Final libraries were visualized on the Agilent 2200 TapeStation fragment analyzer and pooled for 100bp read length single-end sequencing (Illumina HiSeq 2500; University of Wisconsin/Madison).

**Table 2:** Proportions of *Gila robusta* and *G. cypha* assigned genealogically at each sample site. Site=abbreviated locality identifier for each species; P_0_=Pure *robusta*; P_1_=Pure *cypha*; F_1_=First filial hybrid; F_2_=second filial hybrid; B_0_=*G. robusta*-backcrossed hybrid; B_1_=*G. cypha*-backcrossed hybrid; F_N_=late-generation or uncertain hybrid. Samples were assigned to a genealogical class per posterior probability ≥0.80, as assessed using 250,000 post burn-in MCMC generations in NEWHYBRIDS.

| Site | P<sub>0</sub> | P<sub>1</sub> | F<sub>1</sub> | F<sub>2</sub> | B<sub>0</sub> | B<sub>1</sub> | F<sub>N</sub> |
| --- | --- | --- | --- | --- | --- | --- | --- |
| HLCR | - | 1.000 | - | - | - | - | - |
| HGCN | - | 1.000 | - | - | - | - | - |
| HDES | - | 0.083 | - | 0.083 | - | 0.542 | 0.292 |
| HWWC | - | 0.150 | - | 0.200 | - | 0.550 | 0.100 |
| HBKR | - | 0.615 | - | 0.231 | - | 0.077 | 0.077 |
| HYAM | - | - | - | 0.667 | - | 0.333 | - |
| RDES | 0.046 | 0.046 | - | 0.136 | - | 0.364 | 0.409 |
| RSRR ('17) | - | - | - | 0.083 | 0.667 | - | 0.250 |
| RSRR ('09) | - | - | - | 0.455 | 0.455 | - | 0.091 |
| RNJA | 1.000 | - | - | - | - | - | - |
| RMCO | 0.800 | - | - | - | 0.100 | - | 0.100 |
| RWWC | 0.875 | - | - | - | 0.125 | - | 0.375 |
| RC15 | 1.000 | - | - | - | - | - | - |
| RBKR | 0.857 | - | - | - | - | - | 0.143 |
| RECC | 0.938 | - | - | - | 0.063 | - | - |
| RWRW | 0.875 | - | - | - | 0.125 | - | - |
| RUGR | 1.000 | - | - | - | - | - | - |
| RMGR | 0.955 | - | - | - | - | - | 0.045 |
| RYAM | 1.000 | - | - | - | - | - | - |
| RLYC | 1.000 | - | - | - | - | - | - |
| RLSR | 1.000 | - | - | - | - | - | - |

### Assembly and filtering of genomic data

Data assembly was performed using computing resources at the Arkansas High Performance Computing Center (AHPCC), and the XSEDE-funded cloud computing resource JetStream (co-managed by the Pervasive Technology Institute/Indiana University, and the Texas Advanced Computing Center/Austin).

Raw Illumina reads were demultiplexed and filtered using the PYRAD pipeline (Eaton, 2014). Discarded reads exhibited >1 mismatch in the barcode sequence or >5 nucleotides with Phred quality <20. Loci were then clustered *de novo* within and among samples using a distance threshold of 80%. We then removed loci with: >5 ambiguous nucleotides; >10 heterozygous sites in the consensus sequence; >2 haplotypes per individual; <20X and >500X coverage per individual; >70% heterozygosity per-site among individuals; or <50% individual coverage. Individuals with >50% missing data were also discarded. Scripts for post-assembly filtering and file conversion are available as open-source (github.com/tkchafin/scripts).

### Estimating population and individual ancestry

Hypotheses of admixture and hybridization were based on genetic differentiation, as visualized using Discriminate Analysis of Principal Components (DAPC; R-package *adegenet;* Jombart, 2008). Discriminant functions combine principal components (PCs) so as to maximally separate hypothesized groups. Importantly, sufficient PC axes must be retained so as to summarize the high-dimensional input, yet also avoid over-fitting. We accomplished this using the following cross-validation procedure: Stratified random sampling defined 80% of samples per population as a “training set,” with the remaining 20% then classified. PC retention was optimized by minimizing root-mean-square error (RMSE) while maximizing classification success across analyses.

These results were contrasted with model-based assignment tests (STRUCTURE, Pritchard *et al*. 2000; ADMIXTURE, Alexander and Novembre 2009). A shared assumption is that populations can be divided into *K*-clusters identified by permuting membership so as to minimize linkage disequilibrium and departure from Hardy-Weinberg expectations. Given excessive runtimes in STRUCTURE, we first applied ADMIXTURE to evaluate a broader range of models (i.e., *K*=1-20, using 20 replicates), followed by STRUCTURE on a reduced range (*K*=1-10, using 10 replicates with 500,000 MCMC iterations following a burn-in period of 200,000).

Model selection followed a cross-validation procedure in ADMIXTURE where assignment error was minimized by optimal choice of *K*, with results parsed using available pipelines (github.com/mussmann82/admixturePipeline). We used the delta *K* method (Evanno *et al*. 2005) to define the proper model in STRUCTURE (CLUMPAK; Kopelman *et al*. 2015).

We identified putative admixed individuals using Bayesian genealogical assignment (NEWHYBRIDS, Anderson and Thompson 2002) that assessed the posterior probability of assignment to genealogical classes (e.g. F_1_, F_2_), as defined by expected genotype frequency distributions. This component is vital, in that mixed probability of assignment in STRUCTURE and ADMIXTURE can stem from weakly differentiated gene pools. The MCMC procedure in NEWHYBRIDS was run for 4,000,000 iterations following 1,000,000 burn-in, using a panel of 200 loci containing the highest among-population differentiation (*F*_ST_) and lowest linkage disequilibrium (*r*^2^ < 0.2), as calculated in GENEPOPEDIT (Stanley *et al*. 2017). To ensure accuracy of this method as applied to our data, we performed a power analysis using the HYBRIDDETECTIVE workflow (Wringe *et al*. 2017). We first generated simulated multi-generational hybrids using 50% as a training dataset, and then analyzed classification success across replicated simulations using the remaining 50% of samples as a validation set. To examine convergence, simulations were run across three replicates, each with three independent MCMC chains.

### Spatial and genomic heterogeneity in introgression

We tested for signatures of reproductive isolation by examining clinal patterns in locus-specific ancestry across hybrid genomes, using multinomial regression to predict genotypes as a function of genome-wide ancestry. Analyses were performed in the R-package *introgress* (Gompert and Buerkle 2010). Putatively ‘pure’ populations of *G. robusta* and *G. cypha* were diagnosed from results generated by NEWHYBRIDS. We first filtered loci to include those with allele frequencies that differed in the reference populations (as defined by *δ* >0.8, where *δ* is the allele frequency differential at a given locus; Gregorius and Roberds 1986). We generated a null distribution by randomly re-assigning genotypes across 1,000 permutations, so as to test for deviations from neutral expectations. The significance of locus-specific clines (fit via multinomial regression) was then determined by computing a log-likelihood ratio of inferred clinal models versus the null model (at *P*<0.001).

To test for localized breakdown in reproductive barriers, we examined congruence of locus-specific introgression among sampling localities. We did so by deriving site-wise genomic clines within species, then subsequently contrasting the fit of site-wise regression models to the global pattern for each locus. This was accomplished by estimating probabilities of the observed genotypes for each site (*X*_*i,j*_ where *X*=genotypic data over *i* sites for each locus *j*) given the site-specific models (*M*_*i,j*_) versus the range-wide model (*M*_*global,j*_). Concordance was reported as the log-likelihood ratio of *L*(*M*_*global,j*_ | *X*_*i,j*_) to *L*(*M*_*i,j*_ | *X*_*i,j*_) computed per-locus (Gompert and Buerkle 2009).

### Testing effects of anthropogenic pressures

To test correlations between anthropogenic pressures on rates of hybridization, we parsed pressure indices per river reach for four dimensions of human impact from the global stream classifications of Grill *et al*. (2019). These were: 1) River fragmentation (=degree of fragmentation; DOF); 2) Flow regulation (=degree of regulation; DOR); 3) Sediment trapping (=SED); and 4) Water consumption (=USE), from the global stream classifications of Grill *et al*. (2019). We also tested predictive capacity of an integrated multi-criterion connectivity status index (=CSI), also from the free-flowing river assessments of Grill *et al*. (2019). Briefly, the DOF index (from 0 to 100) represents the flow disruption on a reach from dams, while also considering natural barriers such as waterfalls. The DOR index is derived from the relationship between storage volumes of reservoirs and annual river flows and is expressed as the percentage of total river flow that can be withheld in the reservoirs of a river reach. SED and USE quantify the potential sediment load trapped by dams, and the long-term average anthropogenic water consumption as a percentage of natural flow, respectively. The CSI index is a weighted average of these pressure indicators, while also considering road densities and degrees of urbanization [see Grill *et al*. (2019) for details regarding derivation of these indices and their underlying data sources].

We assigned pressure index values for all sites containing at least 1 hybrid (as classified using a 0.90 posterior probability threshold), and tested the predictive power of each pressure dimension on ‘genetic purity’ (calculated via linear regression as the proportion of individuals per population assigned to either P_0_ or P_1_).

## Results

A mean of 106,061 loci were assembled per sample (*σ*=42,689). Following quality/depth filtering, and with mean coverage of 88X, this yielded 16,001 per sample (*σ*=6427). Loci were removed if absent in <50% of individuals, with paralog filtering performed on the basis of allele count and excess heterozygosity. This resulted in 13,538 loci (*μ*=10,202; *σ*=3601), and 1,257,356 nucleotides. Putative orthologs contained 62,552 SNPs, of which 38,750 were parsimony-informative, corresponding to 4.9% and 3% of sampled nucleotides. We retained one SNP per locus, with a final dataset comprising12,478 unlinked SNPs.

### Population structure

Choice of *K* varied by assignment test, with *K*=8 (ADMIXTURE; Fig. S1, S2, and *K*=5 (STRUCTURE; Fig. S2). We thus retained *K*-values from 5-8.

The discriminant function axis with the greatest differentiation (DF1) primarily segregated *G. robusta* in the Little Colorado River (=RECC) from the remaining *G. robusta* and *G. cypha*, with DA2 differentiating Upper Colorado *G. robusta* from *G. cypha* (Fig. 2A). Both assignment tests (Fig. 3) differentiated Little Colorado *G. robusta* from conspecifics, with *G. elegans* forming a discrete cluster. Interestingly, DA3 (Fig. 2B) seemingly identified structure within *G. cypha* as well as potentially admixed populations of *G. robusta*. STRUCTURE models with *K*>8 and ADMIXTURE *K*>5 showed similar restrictions in gene flow between Grand Canyon *G. cypha versus* upper basin sites. Desolation Canyon showed the highest probability of assignment to an ‘upper basin’ *G. cypha* cluster. Within *G. robusta*, a weak signal of reduced intraspecific gene flow was apparent when Green River tributaries were compared with the mainstem Colorado River (i.e. *K*=8, both models).

**Figure 2:**
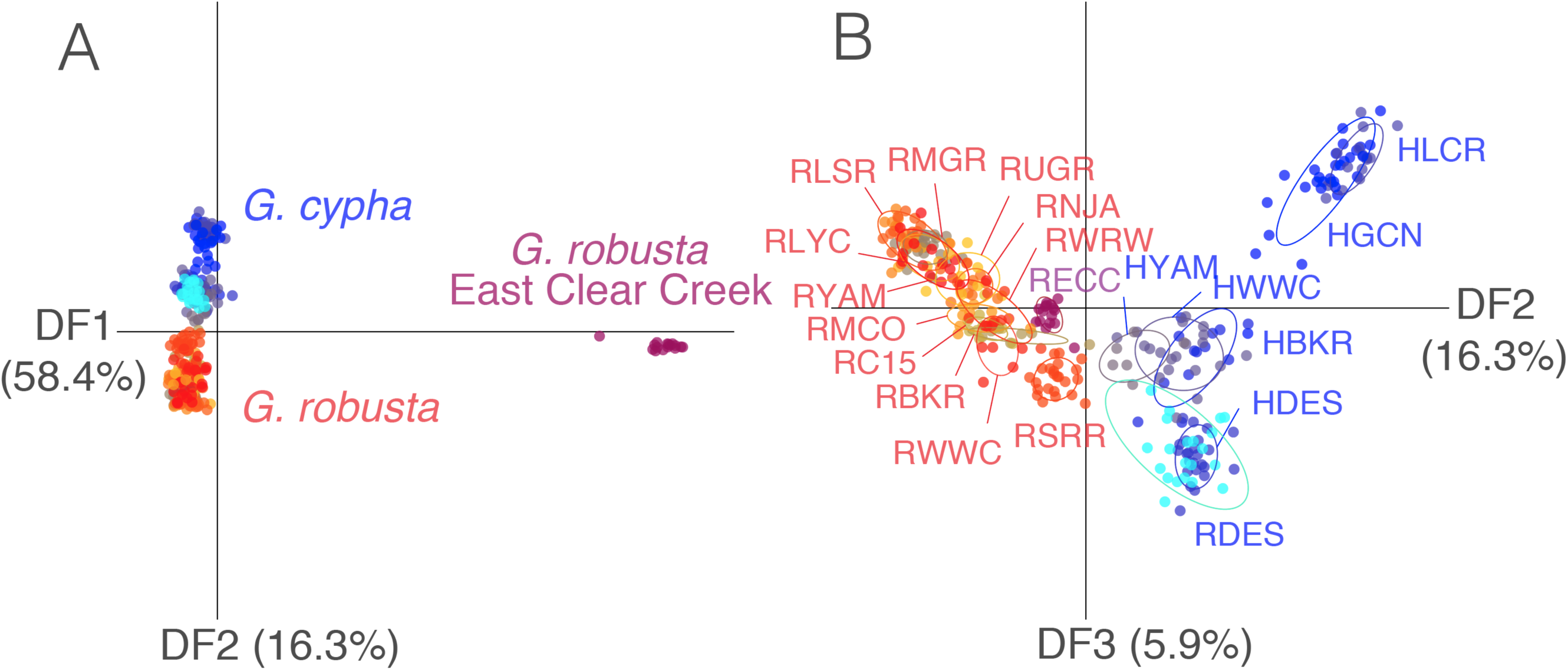
Results of a Discriminate Analysis of Principal Components (DAPC) analysis depicting *Gila robusta* (red), *G. cypha* (blue), and their respective populations (as colored). (A) discriminant function axes 1 and 2 (=DF1xDF2) showing discrimination among both species; (B) axes 2 and 3 (=DF2xDF3) reflecting the manner by which populations of each species (grouped within ellipses) are distributed in discriminant space. The relative percent variance captured by each discriminant function is presented in parentheses. Sample localities are defined in Table 1.

**Figure 3:**
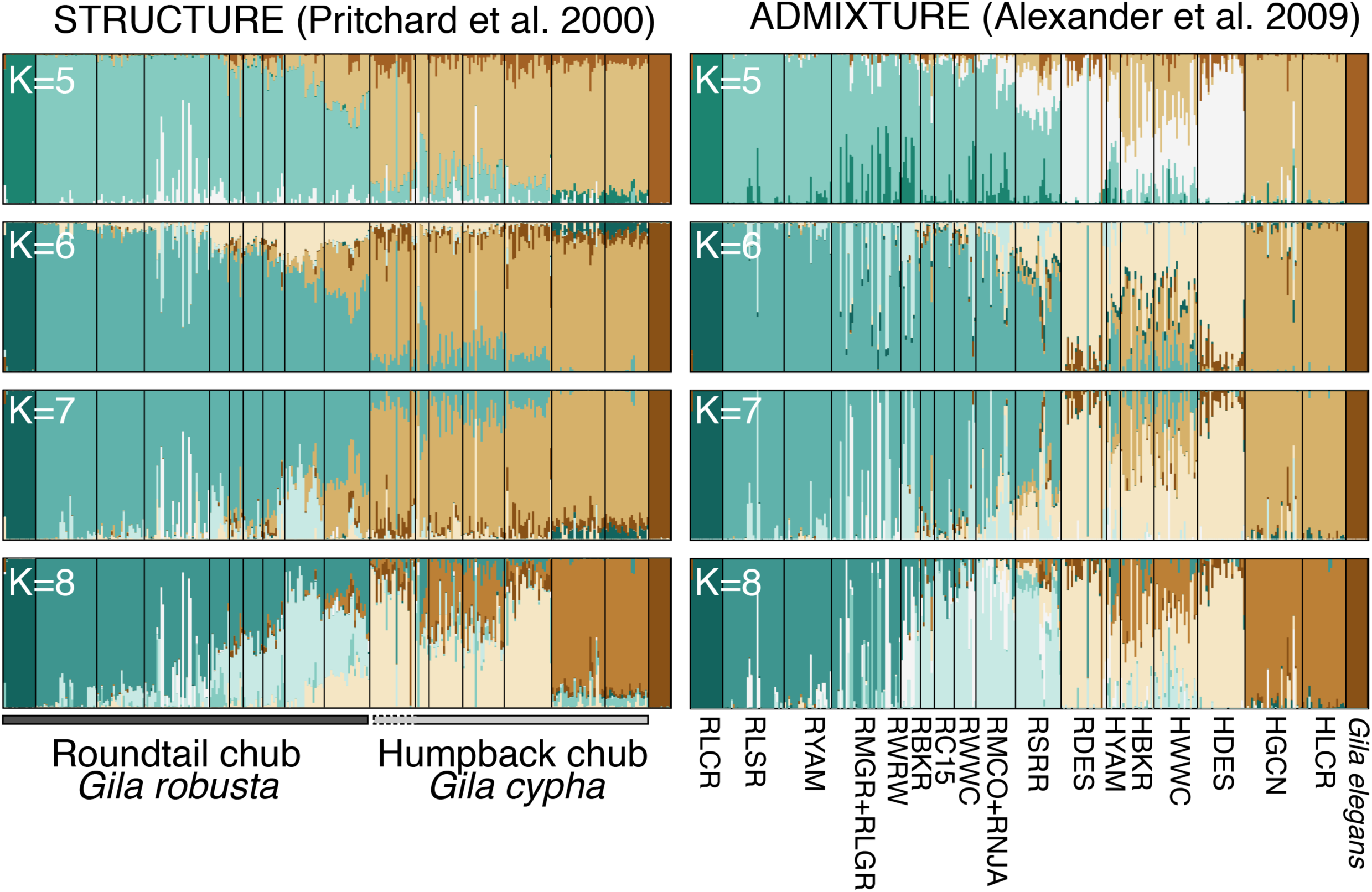
Assignment results for ADMIXTURE and STRUCTURE analyses involving *Gila robusta, G. cypha*, and *G, elegans. K*-values range from STRUCTURE optimum *K*=5 (see Fig. S2) to ADMIXTURE optimum *K*=8 (see Fig. S3). Locality abbreviations are as defined in Table 1.

DAPC and assignment tests each indicated potential hybridization among *G. cypha* and *G. robusta*, most prominently in regions of sympatry (i.e. Black Rocks, Westwater, and Yampa canyons; Fig. 3). Those *G. robusta* sites most ‘distant’ in multivariate space (Fig. 2B) generally showed least probability of interspecific assignment (Fig. 3), presumably due to low or nonexistent admixture with *G. cypha*. Signals of asymmetric introgression were apparent when sympatric localities were examined, with *G. cypha* generally having higher levels of heterospecific assignment.

One exception was Desolation Canyon, where all specimens phenotypically identified as “*G. robusta*” were genetically indistinguishable from those designated as *G. cypha*. Misidentifications at time of capture is a likely cause, owing to the morphological intermediacy of *Gila spp*. at this site (i.e. McElroy and Douglas 1995).

Allopatric populations of *G. robusta* showed less interspecific ancestry, with the exception being the San Rafael River, where samples had 30-50% assignment to *G. cypha* ancestry (Fig. 3), a pattern supported by the weak differentiation of San Rafael in DAPC analyses. Allopatric *G. robusta* from the San Juan River also showed mixed probability of assignment to *G. cypha*, albeit with low probability and consistency.

### Hybrid detection and genealogical assignment

Genealogical assignment in NEWHYBRIDS was used to parse STRUCTURE and ADMIXTURE results for contemporary hybridization. We first defined a prior probability of genetic purity for *G. robusta* as being the upper-most Little Snake River tributaries, and for *G. cypha* as the Little Colorado River confluence in Grand Canyon. Both were chosen based on STRUCTURE and ADMIXTURE results (Fig. 3), and additionally informed by prior studies of natural recruitment (Douglas and Douglas 2010; Kaeding and Zimmerman 1983). As hypothesized, introgressive hybridization at sympatric locations was found to be asymmetric (Fig. 4). Individuals were ‘assigned’ to genealogical class when posterior probabilities exceeded a critical threshold of 0.90 (based upon a <0.025% classification error in power analyses Fig. S4). In cases of mixed assignment, individuals were classified as either “late-generation hybrid,” or “of uncertain status” (Table 2). *Gila robusta* were largely classified as pure in both sympatric and allopatric sites, with a few exceptions (outlined below). The minority of samples assigned to hybrid classes tended to be *robusta*-backcrossed. In contrast, *G. cypha* at sympatric localities had comparatively low purity (0–61%), with most hybrids categorized as either F_2_, *cypha*-backcrossed, or unclassifiable (Table 2). The genetic effects of hybridization are thus inferred as asymmetric, with a greater penetration of introgressed alleles into *G. cypha* populations. For both species, Desolation Canyon samples were mostly classified as either late-generation or unassignable. F_1_ hybrids were notably absent at all localities, suggesting hybridization occurred over multiple generations and ongoing introgression (i.e., hybrids fertile and reproductively successful).

**Figure 4:**
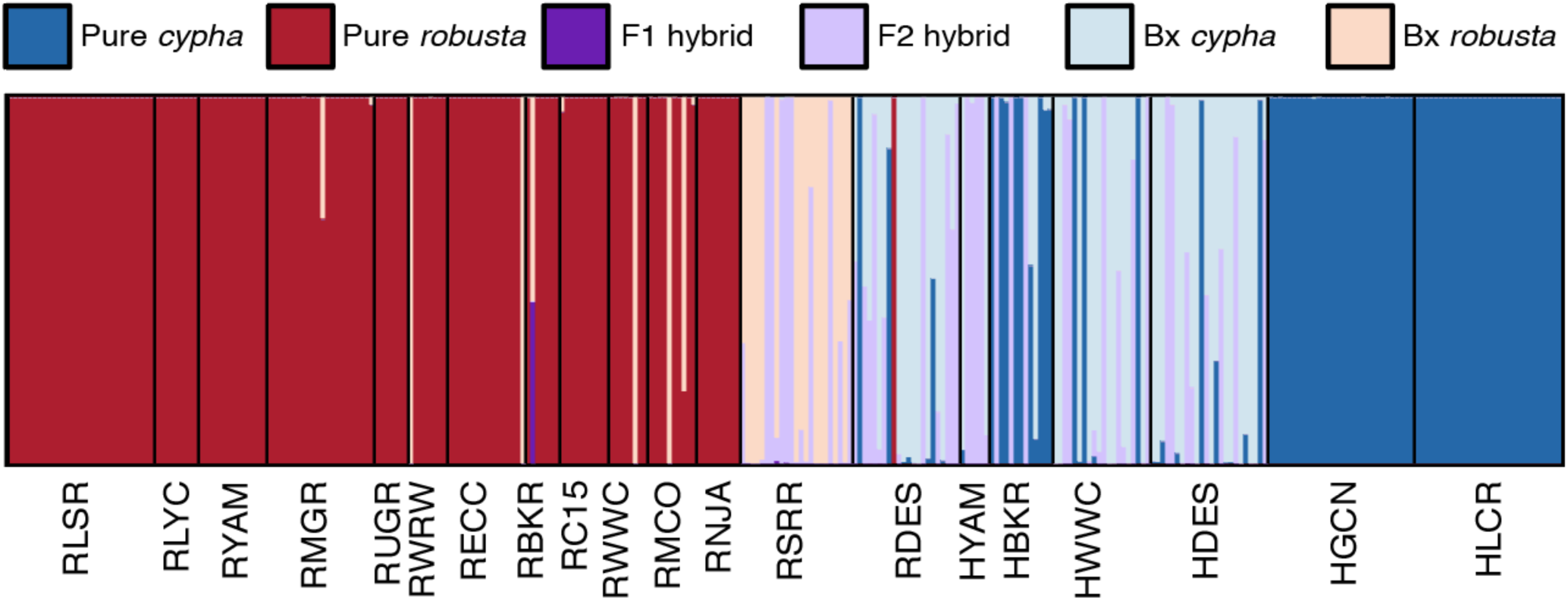
Genealogical assignment for individual *Gila robusta* and *G. cypha*, as compiled from NEWHYBRIDS analysis. Individuals are represented by colored bars, with proportion of color indicating posterior probability of assignment per genealogical class. Prior ‘parental’ allele frequencies for *G. cypha* were derived from the Little Colorado River (HLCR) and from the Little Snake River (RLSR) for *G. robusta* (alternative prior assignments had no significant affect; see Fig. S4 and S5). Colors are as follows: Red=pure *G. robusta*; Blue=pure *G. cypha*; Purple=F_1_ hybrid; Light purple=F_2_ hybrid; Light blue=*cypha*-backcrossed hybrid;Light red=*robusta*-backcrossed hybrid.

Both species showed little signal of contemporary hybridization (i.e., F_1_ hybrids) at allopatric locations, but with notable exceptions being the San Rafael River and, to a lesser extent, the Mancos River. Nearly all San Rafael *G. robusta* were assigned with high probability as either F_2_ or *G. robusta*-backcrossed hybrids, a pattern consistent across years (2009 *versus* 2017), and regardless of priors used. Samples from 2009 were mostly classified as F_2_ (45%) or *robusta-*backcrosses (45%). However, the greatest proportion of 2017 samples were *robusta*-backcrosses (67%) or late-generation hybrids (25%), suggesting an increase of admixture over time (although increased sampling is needed to verify this trend; two-tailed Fisher’s exact test *p*=0.0967; Table 2). The Mancos River samples, composed of 20% hybrids (Table 2), were derived from hatchery stock, not a natural population. Thus, we cannot say if our results represent natural or accidental hybridization that coincided with, or was subsequent to, stock establishment.

### Genomic clines

We also examined how introgression varied across significantly differentiated genomic SNPs and species-diagnostic markers. Here, we considered locus-specific ancestry as the probability of sampling a homozygous *G. cypha* genotype [i.e. P(AA)] as a function of genome-wide ancestry, with the expectation that scant bias should occur if fitness is independent of hybrid ancestry. All loci exhibited clinal patterns that deviated significantly from neutral expectations (*p*>0.001, estimated via permutation; Fig. 5A). The majority displayed coincident sigmoidal relationships between genome-wide ancestry (hybrid index; *h*) and locus-specific ancestry (*ϕ*), although with some divergence. The sigmoidal pattern suggested a deficiency in interspecific heterozygosity, presumably reflecting heterozygote disadvantage (Fitzpatrick, 2013). Notably, some locus-specific clines showed alternative forms (Fig. 5A), indicating that gene flow may be driven by other processes in a minority of genomic regions.

**Figure 5:**
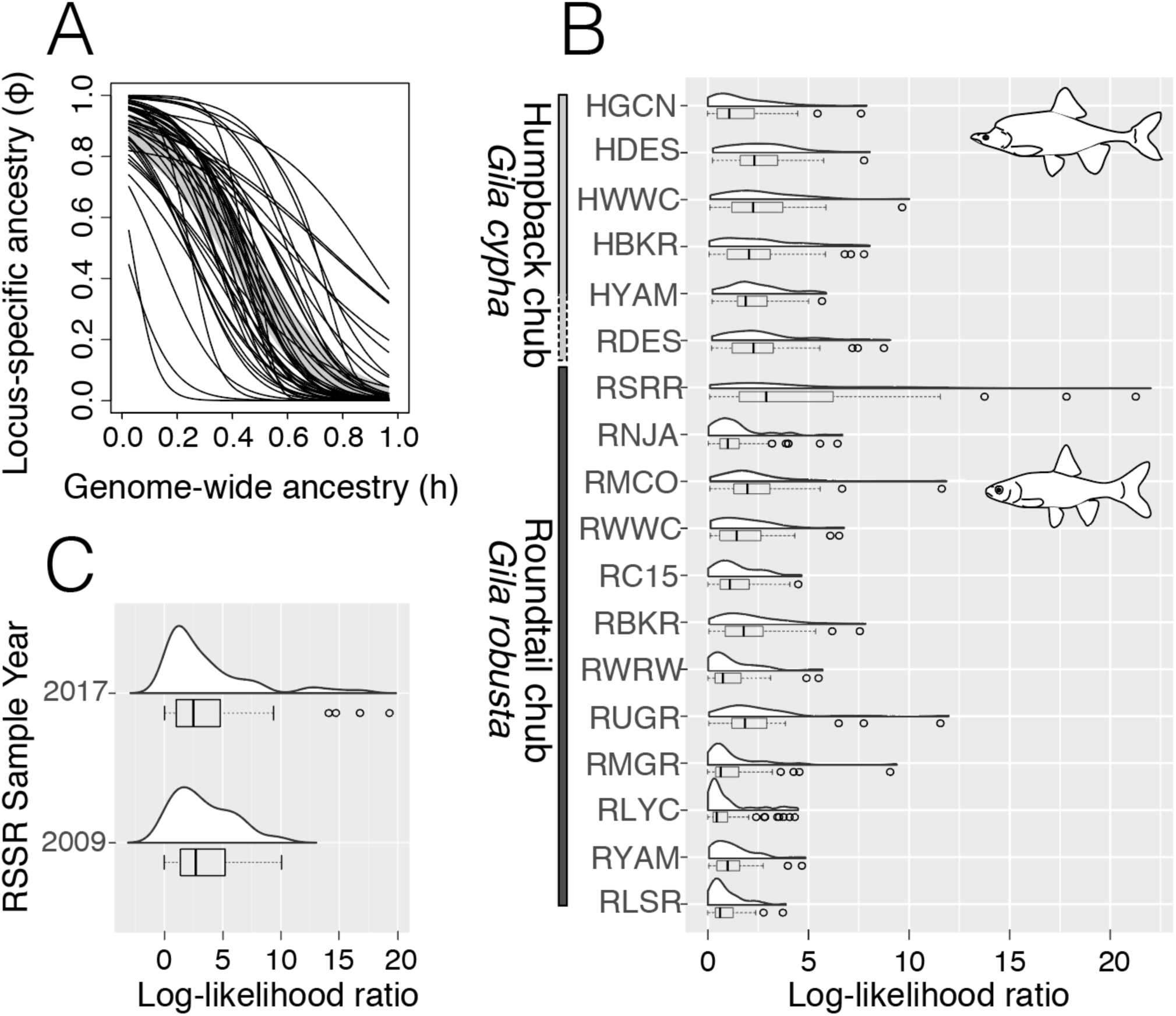
Genomic cline analyses for populations of *Gila robusta* and *G. cypha*, presented as: Per-locus clinal relationships for 50 SNPs with *δ* >0.8 (all significantly non-neutral at *α*=0.001) compared to the neutral expectation (shaded gray region); (B) Log-likelihood ratio distribution of site-wise per-locus clines compared to the global pattern, where higher log-likelihood ratio indicates greater discordance; (C) Per-locus incongruence in genomic clines in the *Gila robusta* samples from the San Rafael River, partitioned by year (2009 versus 2017). Locality codes for populations of each species are defined in Table 1.

We also examined the observed genotypes at each locus, given expectations from the range-wide model and site-specific regression models. These were reported as a log-likelihood ratio *per* locus and *within* each sampling locality (Fig. 5B). We found the ‘fit’ of the range-wide clinal models was rather variable, although with most loci showing little deviation. One notable exception was San Rafael *G. robusta*, where an exceptionally flattened distribution of the locus-347 specific log-likelihood ratios was apparent. This in turn suggested that the global expectation was a poor predictor of within-site genotypes. Thus, while most sites reflected patterns consistent with selection against hybrids, the same cannot be said for the San Rafael River population. It also displayed a strong signal of interspecific admixture in the Bayesian and ML assignment tests (Fig. 3), and variable assignment to >2^nd^ generation hybrid classes in NEWHYBRIDS (Fig. 4), as well as greater intermediacy in multivariate genotypic clustering (Fig. 2). Several other sites also showed a ‘flattened’ distribution of clinal fit among loci (e.g., HWWC, HBKR, HDES, RDES, RUGR, RMCO; see Fig. 5B). This could be a response to elevated introgression and a relative breakdown of heterozygote disadvantage, especially where previous analyses indicated admixture (Fig. 2–4), or as an artefact of reduced sampling (RUGR, RMCO).

Our hypothesis of the San Rafael River as being the most notable discrepancy across all analyses was extended by evaluating samples across different time periods: 2009 (*N*=11) and 2017 (*N*=12). We then fitted locus-specific clines among years (Fig. 5C) and compared those to range-wide expectations (Fig. 5B). We found little change in the overall distribution, save for four outliers in 2017, suggesting further breakdown of clinal expectations over time.

### Testing dimensions of anthropogenic pressure

Among sites showing varying levels of hybridization (N=10; Table 2), only the consumptive water use (=USE) was significantly corelated with a decline in genetic purity (*r*^2^=0.375; *p*=0.035; see Fig. S5). The connectivity status index (CSI) showed a weak but non-significant positive relationship (i.e. increased connectivity = increased genetic purity; *r*^2^=-0.103; *p*=0.7), although we caution that sample sizes were notably low (N=10) after sites were reduced to those containing hybrids and which could be assigned pressure indices (see Fig. S6 for reach assignments).

## Discussion

We found strong evidence for contemporary hybridization among *G. cypha* and *G. robusta* extending beyond their regions of sympatry. These results refine rather than conflict with previous studies employing ‘legacy’ genetic markers (Douglas and Douglas 2007; Dowling and DeMarais 1993; Gerber *et al*. 2001). In addition, these results broaden our understanding of each species and their evolutionary histories, as well as the trajectory of their ongoing evolutionary change in the face of extensive anthropogenic modifications.

### *Species boundaries and reproductive isolation in* Gila

Our survey of the nuclear genome suggested that hybridization between our study species is contemporary in nature, as interpreted from several lines of evidence: The coincidence and shape of our genomic clines; the pervasive signal of genealogical assignment to early-generation hybrid classes; and signatures of selection antagonistic to interspecific heterozygous genotypes.

We interpret these results as remarkable, particularly given the coexistence of study species since at least the mid-Pliocene (Uyeno and Miller, 1965; Spencer *et al*. 2008). In addition, continuing hybridization also stands in stark contrast to the sustained morphological divergence displayed in sympatry (Douglas et al. 1989; McElroy and Douglas 1995). This suggests that genetic exchange is ongoing despite, rather than in the absence of, reproductive isolation.

Dowling and DeMarais (1993) suggested that hybridization between *G. cypha* and *G. robusta* may have contributed to the evolutionary persistence of each species as a distinct entity in the face of historic environmental fluctuations. We concur, and further note that this exchange is ongoing, with a substantial risk that contemporary habitat change will outpace the rate at which introgressed alleles are adaptively “filtered.” If so, then continued habitat alteration could lead to a scenario in which genetic/demographic swamping contributes to local extirpations, or to eventual genetic homogenization of one species by the other (Todesco *et al*. 2016). This pattern is particularly evident in the strong asymmetric signal of gene flow in *G. cypha*, a species of particular concern given its fragmentary distribution and reduced densities within the upper Colorado River Basin (e.g. Badame 2008; Franci *et al*. 2016; USFWS 2017).

To consider the plausibility of such a scenario in which environmental change leads to the dissolution of a species boundary, we must first consider how this boundary is itself structured. We do so by combining the results of our genetic data with those derived from species-specific morphology and life history. Several morphological evaluations have demonstrated that reproductive isolation in extant populations is incomplete (Douglas and Douglas 2007; Douglas *et al*. 2001; McElroy and Douglas 1995). This is likely the result of secondary admixture, rather than a prolonged (i.e., primary) divergence that lead to only weakly-differentiated species. Pliocene fossils that are morphologically consistent with *G. robusta* and *G. cypha* predate major geomorphic and tectonic events that could have triggered vicariant events. For example, the Upper Colorado River was segregated from the contemporary lower basin prior to the mid-Pliocene, (McKee *et al*. 1967), with the uplifting of the Colorado Plateau diverting its flow into one or more Colorado Plateau lakes (Spencer *et al*. 2008). Flows were subsequently diverted by headwater erosion though the Grand Canyon, forming the modern course of the river. Fossil evidence implies that ecological divergence occurred during, or prior to this time, and was sufficient in strength to generate both morphological forms (Uyeno and Miller, 1965). This suggests the existence of ecological conditions that reflect those to which the species are now adapted. Additionally, numerous perturbations [i.e., tectonism, extreme drought (Meko *et al*. 2007)] also occurred during the interval between divergence and present, yet both species not only persisted but did so with some semblance of morphological continuity. Given this, one must again assume that an extant blurring of species-boundaries is a more contemporary occurrence. To test this hypothesis, we considered the ecological dimensions underlying adaptive differentiation in these species.

### *Reproductive barriers in* Gila

Phenotypic and ecological specializations of each species provide potential insights into the mechanisms promoting assortative mating. *Gila cypha* displays phenotypic characteristics interpreted as adaptations to the torrential flows of canyon-bound reaches (McElroy and Douglas 1995; Miller 1946; Valdez and Clemmer 1982). These include a prominent nuchal hump, dorsoventrally flattened head, embedded scales, terete body shape, and a very narrow caudal peduncle that terminates in a caudal fin with a high aspect ratio, indicative of a hydrodynamic shape and powerful propulsion. Its current distribution also reflects association with this type of habitat.

In contrast, *G. robusta* has a comparatively more generalized phenotype, characterized by a deeper and less streamlined body with non-imbedded scales and larger, more falcate fins (Miller 1946). It is found in the upper tributaries of larger rivers (Vanicek and Kramer 1969) with moderate flows. It fails to maintain position within the current when subjected to the extreme flows associated with *G. cypha*, and instead becomes benthic so as to avoid being swept away (Moran *et al*. 2018). This suggests a natural history diametrically opposed to that of *G. cypha*, where dynamic flow regimes clearly predominate. Accordingly, radiotelemetric studies verified habitat preferences for each species, with *G. cypha* seldom straying from the deep eddies and turbulent flows of canyon-bound reaches (Douglas and Marsh 1996; Gerig *et al*. 2014; Kaeding *et al*. 1990). These observations underscore the role that functional morphology plays with regards to species boundaries, in that intermediate morphologies would be maladaptive in either habitat.

However, barriers that sustain reproductive isolation are unclear, in that both species are broadcast-spawners (Johnston and Page 1992), with a temporal overlap in spawning period (Kaeding *et al*. 1990). The latter is likely a consequence of shared environmental cues triggering reproduction, namely seasonal changes in flow rate and temperature, with spatial segregation driven by subsequent alterations in microhabitat and substrate preference (Douglas and Douglas 2000; Minckley 1996). Widespread movements by *G. robusta* during the spawning season contrast with the relative localized focus found in *G. cypha* (Kaeding *et al*. 1990; Tyus *et al*. 1982), and again reinforce the restricted habitat requirements of the latter. Additionally, there is a stronger ‘homing’ component in the microhabitat preferences of *G. cypha* (Valdez and Clemmer 1982). These ecological differences, combined with overall higher abundance of *G. robusta* in most areas (e.g. Francis *et al*. 2015) likely contribute to the observed asymmetric introgression between the two species (Edelaar *et al*. 2008). Intraspecific recognition as a mate-choice mechanism is also an observed behavior that promotes reproductive isolation. Despite congruent reproductive condition and the presence of suitable substrate in a brood stock tank, natural spawning did not occur between *G. robusta* x *G. elegans* and *G. elegans* x *G. cypha* (Hamman 1981).

Thus, we contend that reproductive isolation in *G. robusta* and *G. cypha* is driven by extrinsic factors, with pre-mating isolation primarily in the form of microhabitat selection and post-mating isolation driven by functional morphological differences. Our data point to selection against hybrids as represented by their relatively poor performance in the environment, or to a diminished success in mating. However, we noted a possible breakdown of this expectation at some localities when genomic clines were fitted to within-site patterns. The San Rafael River, for example, is one such exception. Fortney (2015) quantified anthropogenic changes in this river over the last 100 years, with the channel being extensively canalized and diverted, and flows diminished by 83% due to water withdrawals. These manipulations yielded a narrower, relatively deeper channel that stands in sharp contrast to an historically wider and slower river whose flow regime was governed by geomorphology and dominated by flooding. Anthropogenic alterations apparently provided an opportunity for adaptive hybridization, an hypothesis consistent with the exclusive presence of late-generation hybrids in the San Rafael population (Fig. 4). Under this scenario, selective advantage would similarly drive outlier loci and the reduced-fit seen in our clinal models (Fig. 5). The origin of *G. cypha* alleles in this population is unclear, although they may possible be derived from a remnant population in the upper reaches (e.g., Black Box Gorge; Badame, pers. comm).

An examination of the degree to which anthropogenic pressures drive basin-wide hybridization point to a role for consumptive water use in driving a decline in overall genetic purity. Consumptive water usage, and the impact of the associated infrastructures (such as diversions and reservoirs), are often implicated as detrimental to freshwater fish diversity (e.g. Xenopoulos *et al*. 2005). Insofar as river discharge is one dimension of ecological heterogeneity, and given the trend of decreasing species richness as flow declines (Oberdorff *et al*. 1995), we posit that a coincidental relationship is rather extreme in the Colorado River when anthropogenic manipulations and extensive hybridization are contrasted. Yet, a test of this hypothesis is difficult without further experimental work (i.e. leveraging hatchery-produced interspecific hybrids to test for viability in varying habitats, represented within a series of mesocosms). While increased sampling is also necessary, it would be difficult given that we have already sampled 4 of the 5 extant *G. cypha* populations.

### Modified environments and genetic swamping

Grabenstein and Taylor (2018) defined mechanisms that drive anthropogenically-mediated hybridization in coexisting species: 1) Interspecific contact promoted by habitat homogenization or altered phenology; 2) Disruption of mate selection/ choice; and 3) Habitat alteration, such that hybrid genotypes are favored (Anderson 1948). All three are plausible for *Gila*, with the ‘hybrid swarm’ of the San Rafael River an extreme case. Asymmetric hybridization was implicated in all extant sympatric *G. cypha* populations, save the Cataract Canyon aggregate not evaluated in this study. The latter reflects a more ‘*robusta*-like’ morphology (McElroy *et al*. 1997), with low population numbers and a slower growth rate relative to other extant populations (Badame 2008). Taken together, these suggest an elevated risk for genetic or demographic swamping in Cataract Canyon *G. cypha* (Todesco *et al*. 2016), and lend urgency to their inclusion in future genetic surveys.

Such a scenario may also be invoked for *G. cypha* in the Yampa River, recognized even prior to our sampling as being of reduced and declining numbers (Tyus 1998). The ubiquity of highly-admixed genomes in our sampling (from 1999-2001), coupled with the absence of genetically pure individuals in more recent surveys (USFWS 2017), suggest the potential for local extirpation. Given the prevalence of asymmetric hybridization in other sympatric *G. cypha*, it is possible that genetic swamping may have also played a role in the decline of the Yampa Canyon population (although we cannot test that hypothesis). Of note are recent surveys that have documented diminished catch ratios for *G. cypha* at these sympatric localities (Fig. 4; Francis *et al*. 2016; USFWS 2017). Thus, an elevated risk of genetic swamping appears as a strong potential for all *G. cypha* populations sympatric with *G. robusta*.

### Genetic swamping and Allee effects

The capacity for populations to track changing conditions is constrained not only by standing genetic variation but also complex demographic processes that feed back to reproductive fitness (Kokko *et al*. 2018). As the effective population size (*N*_e_) of a population decreases, so also do beneficial variants, primarily due to reduced efficacy of selection relative to genetic drift and associated inbreeding depression (i.e., Allee effects; Kramer *et al*. 2009). This in turn can induce a negative feedback that drives local extirpation (Polechová and Barton 2015). Using a similar logic, we posit that maladaptive introgression within diminishing populations could also synergistically trigger a “runaway” process of genetic swamping (Fig. S7).

In this conceptual model, demographically-driven Allee effects weakens purifying selection against maladaptive introgressed alleles, whereas their continued influx further reduces fitness via outbreeding depression. In this way, maladaptive gene flow can continually depreciate *N*_e_ and effectively promote an “extinction vortex” (Gilpin and Soulé 1986), and we posit this mechanism may contribute to the decline of those *G. cypha* populations sympatric with *G. robusta*. Although some signal of selection against heterospecific alleles was apparent, another manifestation of shrinking *N*_e_ is the expansion of genomic linkage disequilibrium (Nachman 2002). As a result, purifying selection can actually be counterproductive, wherein beneficial genetic variation is lost via selection against linked regions (Nachman and Payseur 2012).

Under this paradigm, the risk of swamping in *G. cypha* is elevated by the numerous factors that increase the relative impact of genetic drift. These are: Reduced population sizes in extant populations (Douglas and Marsh 1996; Tyus 1998); A fragmented distribution (Fagan, 2002); and a “slow” life history (i.e., long generation time and extended lifespans; Olden *et al*. 2008), and higher vulnerability to regulated and reduced flows given its habitat preference of turbulent rivers (as above). The hybrid swarm in the San Rafael River, and the suspected genetic swamping of *G. cypha* in the Yampa River, are potential harbingers of this erosion. Genetic integrity may be preserved in the short term by cultivating “pure” progeny via hatchery production, so as to potentially extend existing pure populations, although a propogation program risks further reducing *N*_e_ (Allendorf *et al*. 2001). The development of pure stock for *G. robusta* should be relatively easy, whereas upper basin *G. cypha* are more problematic in that they display various levels of hybridization (i.e., Figs. 3-4). In this regard, we echo the “producer’s gambit” philosophy (McElroy *et al*. 1997) where hybrid populations fall under an expanded conservation paradigm when genetic purity cannot otherwise be maintained (Lind-Riehl *et al*. 2016). Given apparent ecological non-equivalency of hybrids, we suggest that habitat restoration is the only long-term means to resurrect genetic purity in these populations (Wayne and Shaffer 2016).

### Conclusions

A reduced-representation assay of nuclear genomes in *G. robusta* and *G. cypha* provided evidence of asymmetrical hybridization that is range-wide and spatially heterogeneous (Fig. 3, 4). We interpreted this as reflecting secondary contact, rather than a persistent echo reverberating throughout their respective histories, particularly given the pervasive selection we found with regard to genomic clines operating against interspecific heterozygotes (Fig. 5). Although we lacked appropriate sampling to adequately test for temporal changes in hybridization rates, we did observe the expansion of a hybrid swarm in the San Rafael River over an eight-year period, as well as high levels of asymmetric hybridization in all sympatric populations of *G. cypha* (Table 2). This underscores the potential for genetic/demographic swamping by *G. robusta*, as well as exacerbating the extirpation risk for extant populations of *G. cypha*. We argue that conservation plans for *G. cypha* must consider this possibility. We also suspect the species-boundary for *G. cypha* is largely maintained by extrinsic factors (i.e., lower fitness of hybrid phenotypes and differential microhabitat preferences). As such, further habitat degradation and homogenization may lead to complete genetic erosion, either by contravening habitat selection for pure individuals, or by promoting modified anthropogenic riverscapes that serve as habitat for novel hybrid lineages/swarms. The scenario playing out in *Gila* emphasizes a philosophical dilemma that conservation policy must confront: Is hybridization antagonistic to the conservation of biodiversity or is it instead an adaptive tactic which species employed routinely by species in their evolutionary struggle to persist.

## Acknowledgements

This research was conducted in partial fulfillment by TKC of the Ph.D. degree in Biological Sciences at the University of Arkansas, as enabled by a Distinguished Doctoral Fellowship (DDF) award. Numerous agencies and organizations contributed field expertise, sampling, permit authorization, funding, and/or valuable comments: Arizona Game and Fish Department, Colorado Division of Wildlife, Jicarilla Apache Game and Fish, Nevada Department of Wildlife, United States Fish and Wildlife Service, Utah Department of Natural Resources, Utah Division of Wildlife, and the Wyoming Game and Fish Department. Particular thanks are extended to: J. Alves, M. Anderson, R. Anderson, P. Badame, K. Bestgen, M. Breen, K. Breidinger, S. Bryan, P. Cavalli, B. DeMarais, T. Dowling, R. Fridell, K. Gelwicks, K. Hilwig, M. Hudson, D. Keller, J. Logan, C. McAda, S. Meisuer, T. Modde, K. Morgan, F. Pfeifer, S. Ross, R. Timmons, P. Unmack, D. Weedman, K. Wilson, and E. Woodhouse (with apologies to anyone inadvertently overlooked). Additional sampling was completed by MED and MRD under permits provided by: Arizona Game and Fish Department, Dinosaur National Monument, Grand Canyon National Park/Glen Canyon National Recreation Area, The Hualapai Tribe, The Navajo Nation, Nevada Division of Wildlife, U.S. Fish and Wildlife, and Wyoming Department of Game and Fish. Sampling procedures were approved under Arizona State University Animal Care and Use Committee (ASU IACUC) permit 98–456R and Colorado State University Animal Care and Use Committee (CSU IACUC) permit 01–036A-01. Travel in the Grand Canyon was conducted under auspices of a Grand Canyon National Park river use permit. We are also indebted to students, postdoctorals, and faculty who have promoted our research: A. Alverson, W. Anthonysamy, M. Bangs, M. Davis, L. James, S. Mussmann, J. Pummill, A. Tucker. Funding was provided by several generous endowments from the University of Arkansas: The Bruker Professorship in Life Sciences (MRD), and the Twenty-First Century Chair in Global Change Biology (MED). Additional analytical resources were provided by the Arkansas Economic Development Commission (Arkansas Settlement Proceeds Act of 2000) and the Arkansas High Performance Computing Center (AHPCC), and from an NSF-XSEDE Research Allocation (TG-BIO160065) to access the Jetstream cloud.

## Competing Interests

The authors declare no conflict of interest.

## Data Accessibility

Pending acceptance, all raw sequence files will be uploaded to the NCBI GenBank Short-read Archive (SRA). Relevant curated (i.e. assembled and filtered) datasets will be archived on Dryad. Codes and custom scripts developed in support of this work are available as open-source under the GNU Public License via GitHub: github.com/tkchafin

- Raw sequence files: NCBI SRA (pending)

- Sampling locality metadata, curated sequence files, and alignments: Dryad (pending)

- Codes and scripts: github.com/tkchafin (and as cited in-text)

## Author Contributions

TKC, MED, and MRD conceptualized and designed the research. MED and MRD contributed sampling and coordination of agency efforts. TKC performed molecular work and analyses. TKC and BTM wrote software and codes to analyze data in support of this research. All authors contributed to writing the manuscript and approve of the final submission.

## Supplemental Material

**Figure S1:**
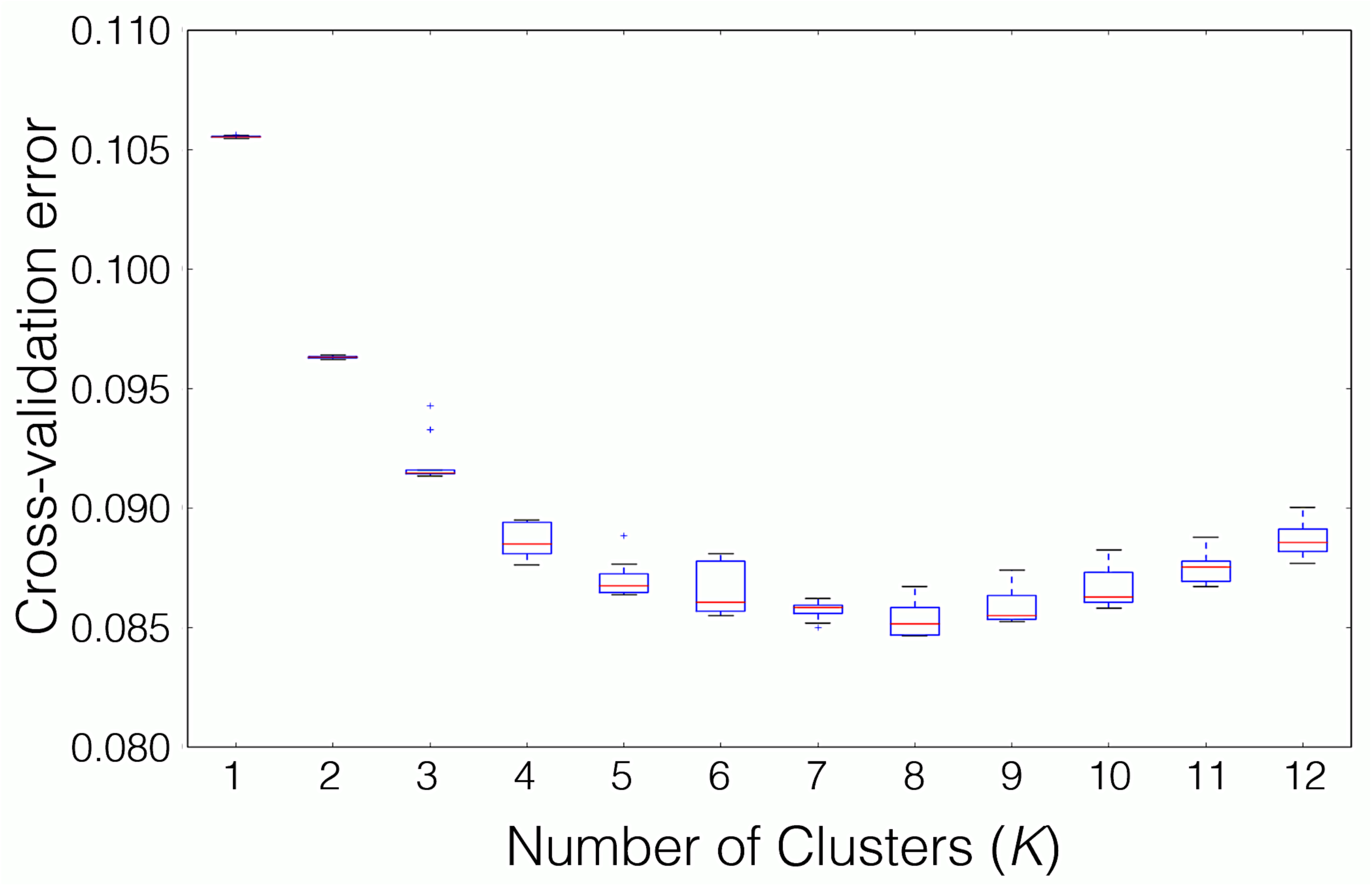
Cross-validation error analysis for ADMIXTURE, as represented by number of clusters (*K*)

**Figure S2:**
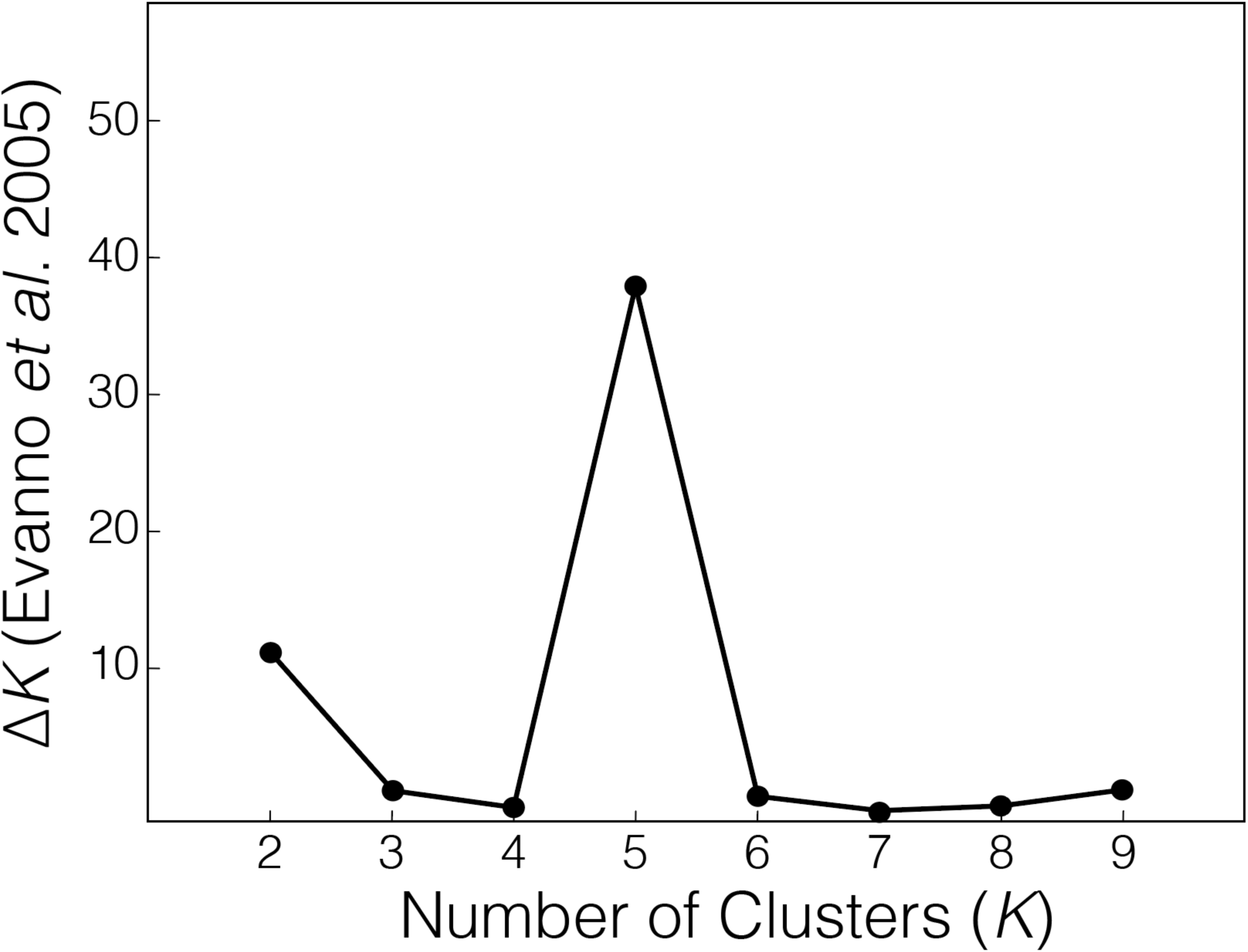
Change in model likelihood by *K* derived from a STRUCTURE analysis.

**Figure S3:**
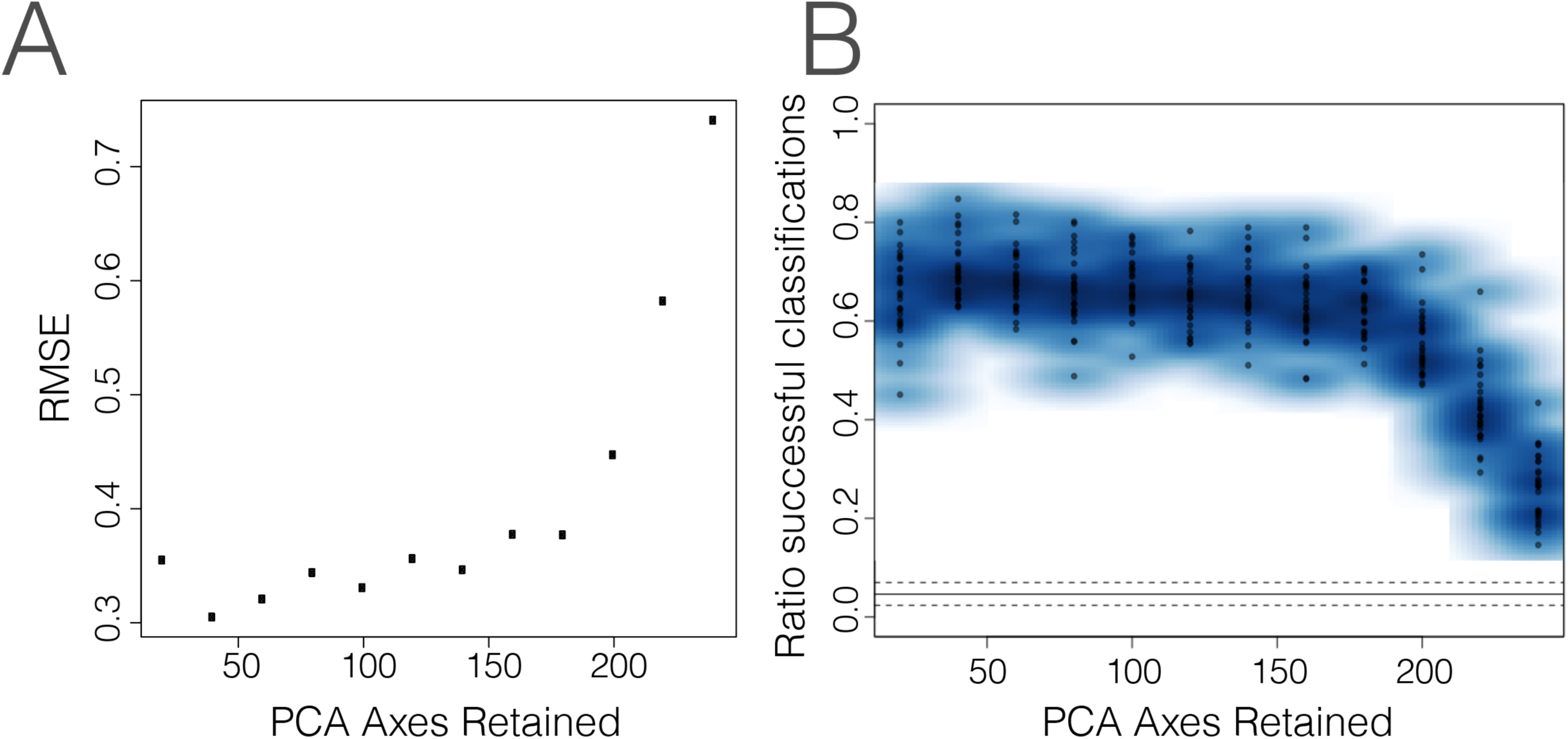
Results of cross-validation analysis in Discriminant Analysis of Principal Components (DAPC), depicting (A) root-mean-square error (RMSE) for classifications under varying number of Principal Component axes (PC’s) retained, and (B) proportion of successful classifications for 20 replicates with varying number of PC’s retained.

**Figure S4:**
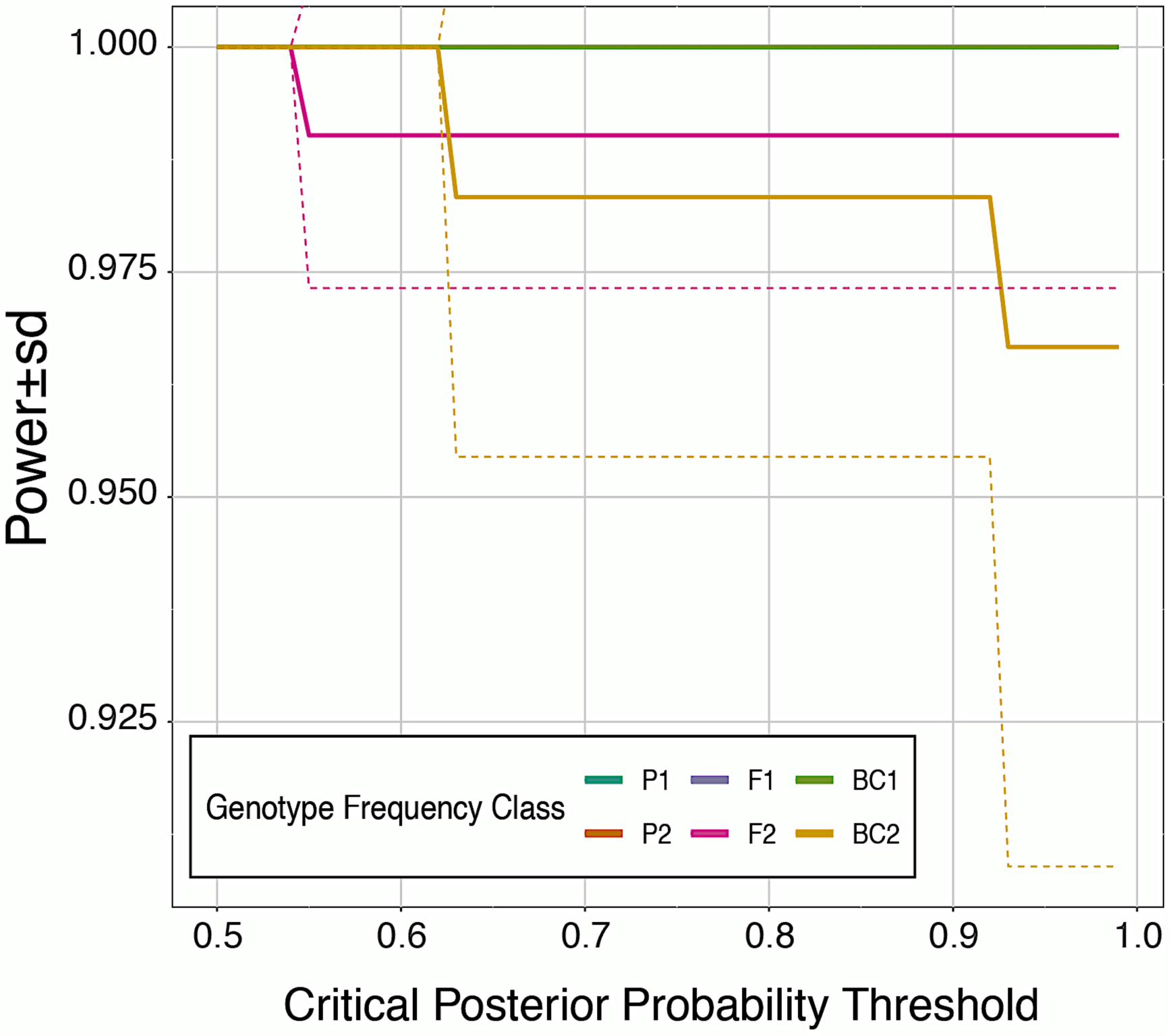
Plot of statistical power of assignment for each of six genotype frequency classes. NewHybrids of simulated hybrid genotypes in HybridDetective, using various critical thresholds (from 0.5 to 1.0) for posterior assignment probabilities. Solid lines indicate mean power, while dashed lines are standard deviations across replicates. Note that accuracy of assignment is 100% in all cases (not shown).

**Figure S5:**
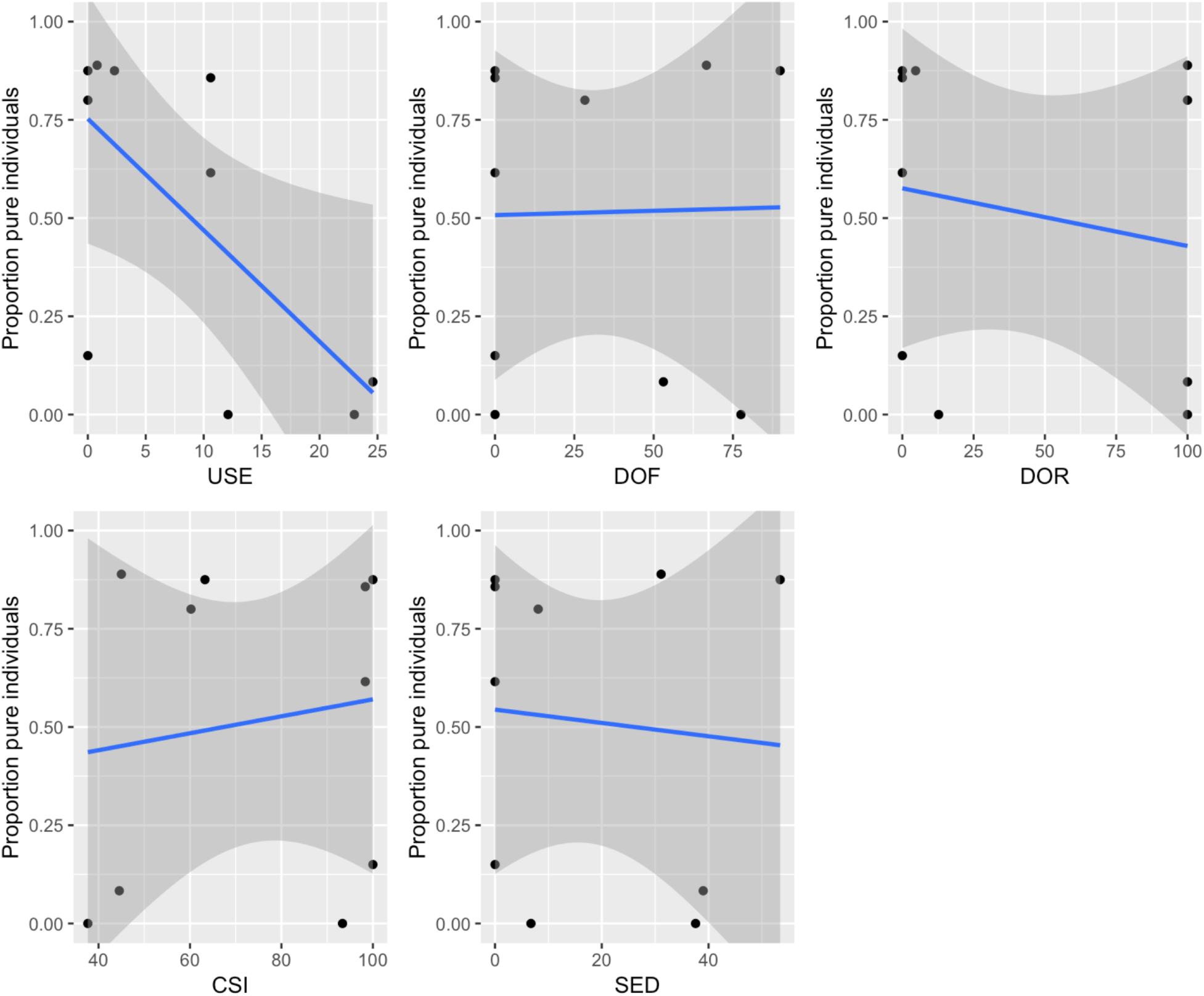
Relationships between five indices of anthropogenic pressure on stream reaches, and the proportion of genetically pure individuals sampled therein. DOF=Degree of fragmentation; DOR=degree of regulation; SED=degree of sediment trapping; USE=percent consumptive water use; CSI=connectivity status index. Scales for DOF, DOR, SED, and USE reflect a proportion of effect, with 100 = 100% impact. CSI scales from 0 to 100, with 100 being full connectivity (i.e. not detectable human impact).

**Figure S6:**
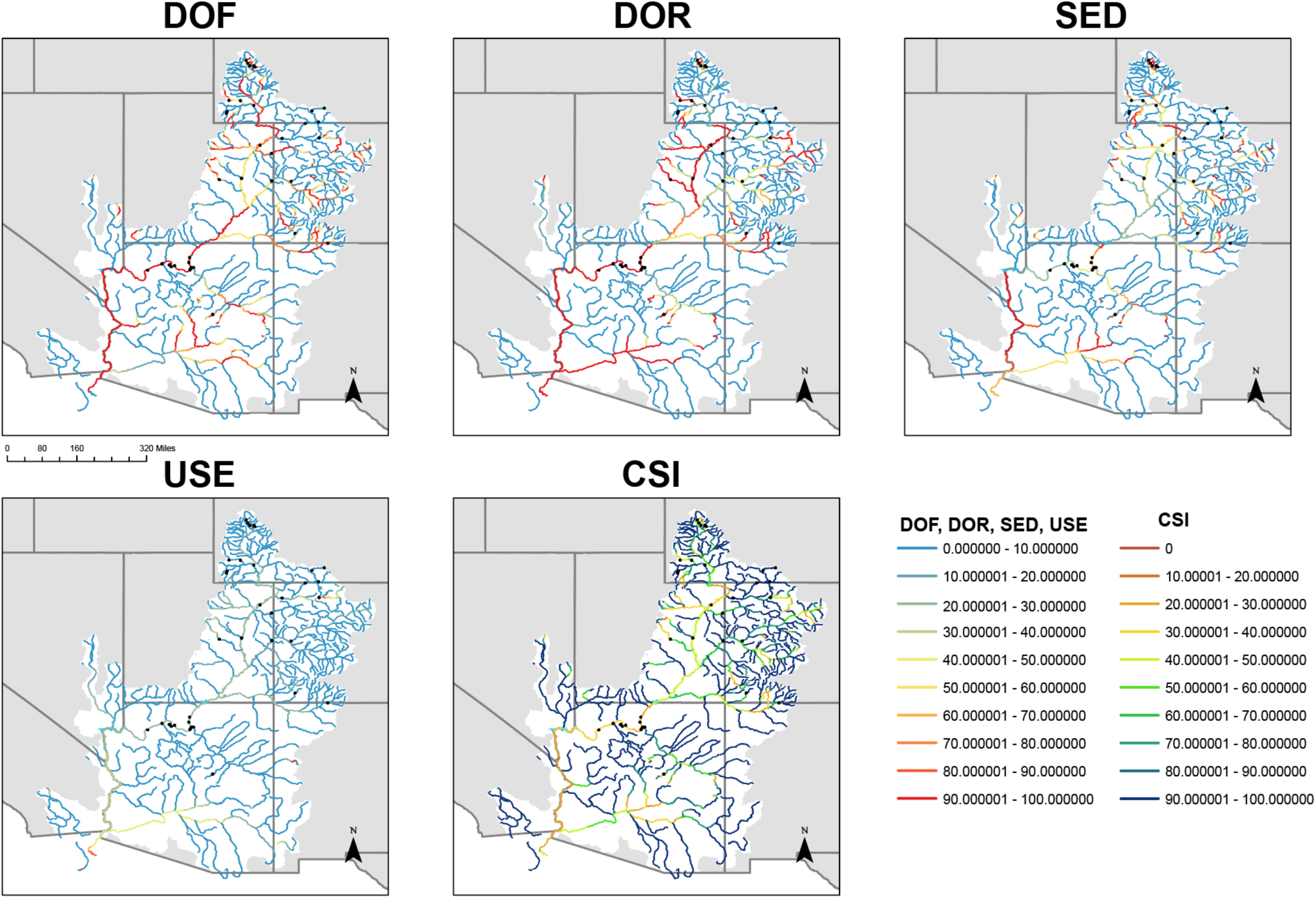
Human pressure indices plotted onto stream reaches in the Colorado River. DOF=Degree of fragmentation; DOR=degree of regulation; SED=degree of sediment trapping; USE=percent consumptive water use; CSI=connectivity status index. Scales for DOF, DOR, SED, and USE reflect a proportion of effect, with 100 = 100% impact. CSI scales from 0 to 100, with 100 being full connectivity (i.e. not detectable human impact).

**Figure S7:**
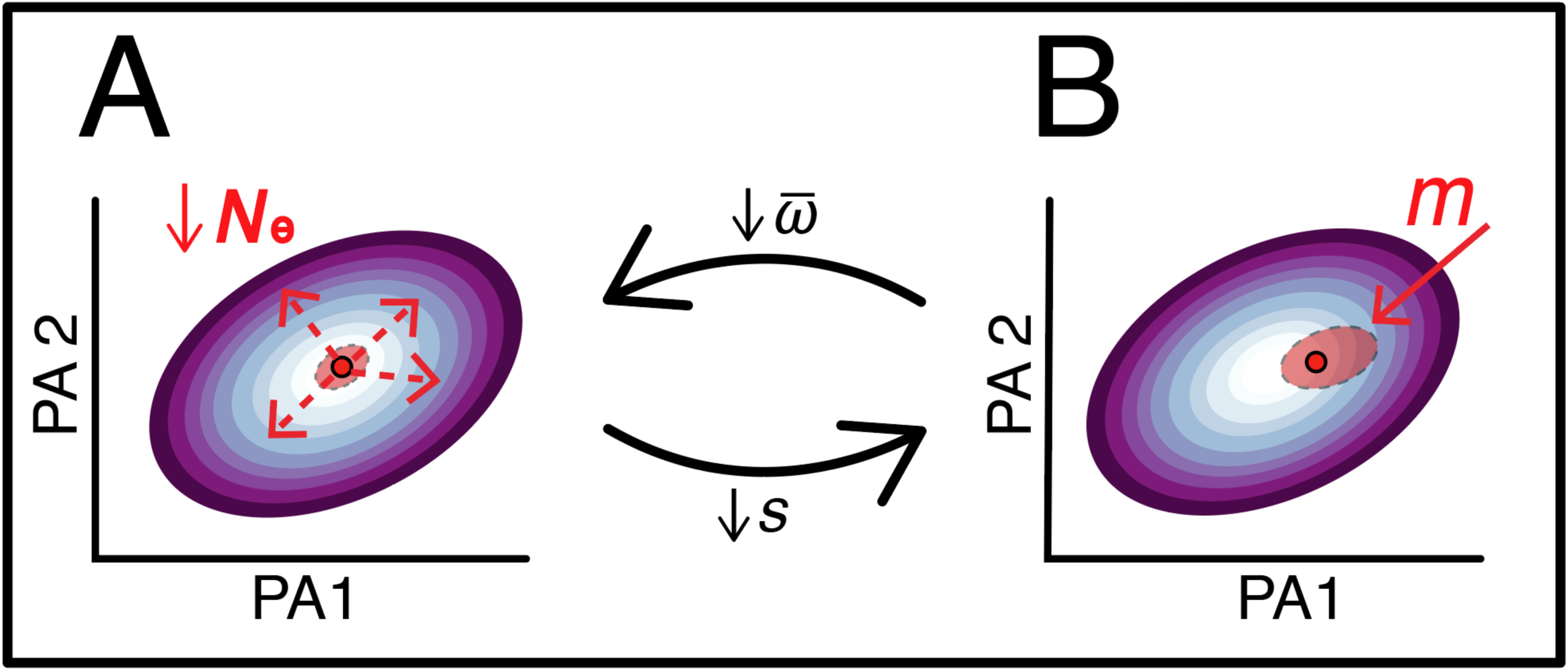
Extinction vortex via negative feedback of inbreeding and outbreeding depression. Shown is the population mean fitness (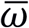; red dot) and variance (red ellipse) within a fitness gradient from low (purple) to high (white), as a function of two arbitrary phenotypic axes (PA1, PA2). (A) Decreasing effective populations size (*N*_e_) lowers the strength of selection (*s*) and reduces both 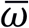 and genetic variance (=inbreeding depression). (B) When introgression is maladaptive, increased gene flow (*m*) bolsters maladaptive genetic variance while driving a further reduction in 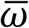 (=outbreeding depression). This, in turn, prompts further depreciation in *N*_e_, with weakened purifying selection and a relatively greater influence of maladaptive *m* as a consequence. Persistent coupling of these processes can then drive population extirpation, especially given extrinsic effects on 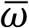, such as rapid environmental change.

